# Motor cortex perineuronal net modulation improves motor function in a Parkinson’s disease mouse model

**DOI:** 10.1101/2024.05.31.596849

**Authors:** David Benacom, Camille Chataing, Alain Prochiantz, Ariel A. Di Nardo

## Abstract

The 6-OHDA mouse model recapitulates midbrain dopaminergic cell loss and associated motor deficits akin to those observed in Parkinson’s disease. Emerging evidence suggests that modulating interneurons in the primary motor cortex could offer a means to mitigate symptoms. In the cortex, perineuronal nets (PNNs), a specialized extracellular matrix structure generally present around fast-spiking parvalbumin interneurons, can modulate neural activity and circuit plasticity. We found that removing PNNs through unilateral or bilateral ChABC injection in the motor cortex temporarily altered motor behavior. Surprisingly, bilateral reduced motor cortex PNNs are observed two weeks after unilateral 6-OHDA midbrain lesions, whereas five weeks after lesion, PNNs return to control levels. Subsequent bilateral ChABC injections significantly improved motor function in 6-OHDA animals only when associated with motor stimulation involving enriched housing and daily motor training. Thus, PNN modulation in the motor cortex of a Parkinson’s disease model enables local circuits to adapt to the loss of dopaminergic inputs, resulting in improved motor behavior.

## Introduction

Perineuronal nets (PNNs) are specialized extracellular matrix structures that form around specific neurons in the brain and spinal cord (1). They are primarily composed of hyaluronan, chondroitin sulfate proteoglycans (CSPGs), link proteins, and tenascin-R (2, 3). In the cerebral cortex, PNNs are predominantly found around the soma and proximal neurites of fast-spiking parvalbumin-expressing interneurons (PV cells) that propagate gamma oscillations and provide cortical coherence (4, 5). The postnatal maturation of cortical PV cells drives critical periods (CPs) of heightened plasticity that allow neural circuits to be remodeled by experience (6). While their maturation kinetics depends on cortical region, the process is consistently accompanied by the formation of PNNs which contribute to CP closure by limiting synapse turn-over and by binding to signaling factors functioning as plasticity brakes (7–11).

The removal of cortical PNNs reverses PV cell maturation and enhances adult plasticity, offering the potential for brain repair (12). Local parenchymal injections of chondroitinase ABC (ChABC) enzyme degrade CSPGs resulting in rapid yet transient PNN removal, with PNN recovery beginning by 6 days and reaching at least 80% by 3 weeks (7, 13, 14). In rodents, ChABC has been used in the primary visual cortex (V1) to restore binocular vision in models of amblyopia (15), and in the spinal cord and peripheral nerves to treat injuries via its action on glial scars (16–18). Simultaneous ChABC injections in substantia nigra pars compacta (SNc) and striatum protects from the loss of midbrain dopaminergic (mDA) neurons and fibers after partial lesions of the nigrostriatal tract (19). In otherwise healthy adult rodents, PNN removal can impact PV cell function and animal behavior (20–23). For example, ChABC injection in V1 coupled with monocular deprivation can induce amblyopia (12), while injection in basolateral amygdala can erase acquired fear memories (24). In disease or brain injury models, ChABC treatment provides positive behavioral outcomes in rodent models of pathologies such as stroke and Alzheimer’s disease (25–29).

The role of PNNs in the primary motor cortex (M1) of healthy adult mice and Parkinson’s disease (PD) mouse models has not been investigated. PNNs accumulate around M1 PV cells across species (1, 30), and while M1 PV cell activity may not be altered in PD mouse models involving acute 6-hydroxydopamine (6-OHDA) midbrain lesions (31, 32), there is evidence for PV cell activity in alleviating motor symptoms (31). We find that PNN removal in M1 with ChABC treatment can alter motor behavior, and that PNNs are transiently reduced in 6-OHDA lesioned mice. Subsequent bilateral ChABC injections in lesioned animals revealed that PNN removal, coupled with motor stimulation, improved motor performance, indicating that targeted PNN disruption could ameliorate motor deficits in neurodegenerative conditions. This study reveals a nuanced role of M1 PNNs in motor function and recovery, highlighting their potential as therapeutic targets in motor rehabilitation.

## Materials and methods

### Animal management

All animal housing and experimental procedures were carried out in accordance with the recommendations of the European Economic Community (2010/63/UE) and the French National Committee (2013/118). Experiments were performed with female C57BL/6J mice (Janvier) housed with *ad libitum* access to food and water under a 12h light/dark cycle. For surgical procedures, animals were anesthetized with ketamine-xylazine (100 mg/kg Imalgene 1000, Boehringer Ingelheim; 5 mg/kg Rompun 2%, Bayer) by intraperitoneal injection. One week prior to undergoing brain lesions, mice were provided DietGel and Hydrogel *ad libitum* in order to limit weight loss.

### Midbrain lesions

In order to protect noradrenergic neurons from 6-OHDA, animals received an intraperitoneal injection of Desipramine-HCl (20 mg/kg, 3067 Tocris) 20 minutes before anesthesia. Stereotaxic injection of 6-OHDA-HCl (H4381 Sigma, 7.2 µg/µl in 0.02% ascorbate-saline) targeted the medial forebrain bundle (MFB) with 30G flat needle at the coordinates relative to bregma: ML, 1.150 mm; AP, −1.150 mm; DV, −5 mm. The needle was left in position for 2 min prior to the injection of 6-OHDA (0.35 µl for 2.5 µg dose) at a rate of 0.1 µl/min. The needle was left in place for 3 min and then slowly retracted at a rate of 2.5 mm/min. The injections of vehicle solution (0.02% ascorbate-saline) were performed with the same procedure.

After surgery, 500 µL of 5% glucose in saline solution was injected subcutaneously and mice were placed in cages under infrared light (50 W) such that half of the cage remained at ∼35 °C during the first week of recovery. Weight was monitored every second day, and mice were provided DietGel and HydroGel daily to minimize weight loss. Animals presenting signs of distress associated with a >20% weight loss were euthanatized.

### ChABC injections

Stereotaxic injection of recombinant ChABC (50 U/ml, 6877-GH Bio-Techne) was performed with a 33G beveled needle in two distinct sites of M1 at the following coordinates relative to bregma: anterior injection with bevel directed posteriorly, ML, ±1.6 mm; AP, +2.2 mm; DV, −0.5 mm; posterior injection with bevel directed anteriorly, ML, ±1.6 mm; AP, +1.4 mm; DV, −0.8 mm. The needle was left in place for 2 min prior to injection of 0.5 µl of ChABC saline solution at a rate of 0.1 µl/min. The needle was left in place for 3 min and slowly retracted at a rate of 2.5 mm/min. Vehicle injections (saline, 0.9% NaCl) were performed with the same procedure.

### Animal behavior

#### Motor enrichment

Mice undergoing a motor training procedure were housed with regular nesting material, an opaque plastic cylinder (10 cm, ⌀ 4 cm), and a wheel (Innodome™ & Innowheel™, Innovive). Control mice were housed with nesting material only.

#### Rotation test

Rotation behavior was quantified with a rotameter (Ugo Basile). Mice were tested 2 weeks after 6-OHDA lesions. Full 360° rotations were recorded during 5 min either before (spontaneous) or after injection of amphetamine (5 mg/kg), when specified. For recording purposes, a magnet was attached to the tail of the mice before placement in the rotameter. Net ipsilateral rotations were reported as total right minus total left 360° turns.

#### Rotarod

A high-height rotarod apparatus (LSI Letica instruments; LE8200) with the rod situated 50 cm above the bench was used in order to provide animals further motivation to perform the task. To minimize stress upon falling, a platform was inserted 20 cm below the rod for the first trials. For each trial, the time spent by each animal on the rod before falling (latency to fall) was measured.

For healthy animals injected with ChABC or vehicle, daily trials were performed with an accelerating rotarod on “gentle” settings (4 to 40 rpm in 5 min) for 3 days prior to surgery (Test 1), and then for 5 consecutive days starting 1 day after surgery (Test 2) and 17 days after surgery (Test 3). At the end of the first week and at the end of the third week, animals were submitted to a single test on an accelerating rotarod on “brisk” settings (4 to 40 rpm in 2min). For lesioned animals, a pre-screen was performed 4 weeks after 6-OHDA injections in order to validate lesion efficiency and also provide habituation. During the pre-screen, mice were tested once a day, for three consecutive days, at 12 rpm fixed speed. The exclusion criterion was >25 sec before falling, which corresponds to the ability of the worst-performing control animals at this timepoint. Lesioned animals surpassing this threshold were excluded from subsequent experimentation. For every subsequent rotarod session, mice underwent three daily rotarod tests for five consecutive days at 12 rpm fixed speed. Final performance was calculated by taking the average of each day’s best performance.

#### Irregular beam test

The irregular beam consisted of a piece of hardwood (5 cm × 50 cm) with irregularly spaced slots suspended 50 cm above the ground between a platform and the home cage. For habituation, mice were trained to walk the length of the beam twice with gentle pushes using a small wooden cylinder. For the final trial, mice were filmed in order to count the total number of paw slips on both sides of the beam.

#### Pole test

The pole test consisted of a rod placed vertically in the center of the home cage in order to minimize stress. For healthy animals injected with ChABC or vehicle, a metallic pole (30 cm, ⌀ 1 cm) was used in order to maximize the level of difficulty. For all other experiments that include lesioned animals, a wooden pole (30 cm, ⌀ 1 cm) was used. Mice were placed at the top of the pole facing upwards, and the time taken to go from the top of the pole down into the home cage was measured. The ability/accuracy to control the descent was evaluated by the experimenter blind to the mouse group. Motor performance was taken as the mean of 3 tests each scored from 0 (fail) to 5 (best): 5, controlled descent, with the tail around the pole, in less than 30 seconds; 4, uncontrolled descent (animal on its side, or without tail control) in less than 30 seconds; 3, controlled descent, with the tail around the pole, taking more than 30 seconds; 2, uncontrolled descent taking more than 30 seconds; 1, failure to descend (jump or fall) in less than 1 minute; 0, failure to descend after more than 1 minute of freezing at the top of the pole.

#### T-maze test

For the alternate-choice T-maze test, a cross-maze with four equal arms in length (37 cm × 7 cm) was used with one arm closed by a mobile separator to form a T. For habituation, mice were left to explore the maze for 15 min before testing. Thirty trials were performed per mouse with a 30 sec resting period between each trial while the mobile separator was moved in the clockwise direction or anti-clockwise direction to minimize visual cue and habituation effects. The percentage of right-hand turns (turns towards the direction ipsilateral to ChABC injection side) was calculated.

### Tissue collection

For immunohistochemistry (IHC), mice were deeply anesthetized with Ketamine-Xylazine (100 mg/kg Imalgene 1000, Boechringen Ingelheim; 5 mg/kg Rompun 2%, Bayer) and transcardially perfused with PBS followed by 2% PFA in PBS using infusion pumps (KD Scientific). Perfused brains were incubated in 2% PFA at 4 °C overnight, rinsed in PBS and placed in 30% sucrose in PBS at 4 °C until equilibrium before embedding in cryoprotectant (OCT) at ∼-50 °C.

### Immunohistochemistry

Embedded brains were sectioned at 40 μm on a cryostat (HM 560, ThermoScientific) and stored at −20 °C. Brain sections were permeabilized with PBS, 1% Triton (Sigma-Aldrich) for 45 min at RT. Sections were blocked in PBS, 10% normal goat serum (NGS; Thermo Fisher Scientific) for 30 min at RT, incubated with primary antibodies diluted in PBS 1% Triton, 3% NGS overnight at 4 °C, washed 5 times in PBS, 0.1% Triton, incubated in PBS, 3% NGS, 0.1% Triton with secondary antibodies for 2h at RT, washed 5 times in PBS and mounted in Fluoromount-G (Southern Biotech). Primary antibodies were used at the following dilution: PV 1:2000 (Rabbit, Swant 27); WFA 1:500 (Biotin, L1516, Sigma); TH 1:800 (Chicken, Sigma). Corresponding Alexa-fluor secondary antibodies (Thermofisher) were used at a 1:2000 dilution.

### Image analysis

Images were acquired using an Axiozoom or an Axioscan microscope (Zeiss) with ZEN software (Zeiss). Images were captured at 20X magnification using fixed exposure times and light intensities across experiments. For quantifying PV^+^, WFA^+^, and PV^+^WFA^+^ cell numbers in the M1 cortex, the mean cell density per area was measured from 3 sections in the anterior M1 and 3 sections in the posterior M1 for each mouse. Image analysis was performed with Qupath software, with the experimenter blind to experimental group. For TH quantifications, a minimum of 3 sections encompassing the SNc was used to determine the average the number of TH^+^ cells in both lesioned and non-lesioned hemispheres for each mouse.

### Statistical analysis

Statistical analyses were performed with GraphPad Prism (v10.1.1). The *t*-test was used for single comparisons between two groups, while main effects and interactions with more than two groups were determined using analyses of variance (ANOVA) or 2-way ANOVA with Tukey’s test, uncorrected Fisher’s LSD, or Holm-Šídák’s multiple comparison post hoc test. Statistical significance was accepted with *P* < 0.05. Data are expressed as mean ± SD and relevant tests are indicated in the Figure legends.

## Results

### Acute removal of PNNs transiently perturbs motor behavior

To test whether the local removal of PNNs in M1 can affect motor behavior, ChABC was injected unilaterally in a cohort of healthy adult mice (Fig. 1A). This treatment resulted in ∼50% loss of PNNs one day later followed by complete recovery of PNNs by 3 weeks post-injection (wpi) (Fig. 1B). Potential asymmetry in motor response was evaluated by spontaneous and amphetamine-induced rotation tests after 1 day. While mice showed no change in spontaneous motor behavior (Fig. 1C), amphetamine treatment induced significant rotation behavior in the direction contralateral to the side of the ChABC injection (Fig. 1D). The injection of ChABC did not affect the total mean number of spontaneous or amphetamine-induced rotations (Figs. S1A and S1B). The lateralization of amphetamine-induced motor behavior was confirmed by the alternate-choice T-maze test, in which ChABC-treated mice turned significantly in the contralateral direction (Fig. 1E). After 3 weeks, amphetamine-induced rotation behavior was no longer observed in either test (Figs. 1D and 1E), which is in keeping with the recovery of PNNs (Fig. 1B) and suggests effects are transient.

**Figure 1.**
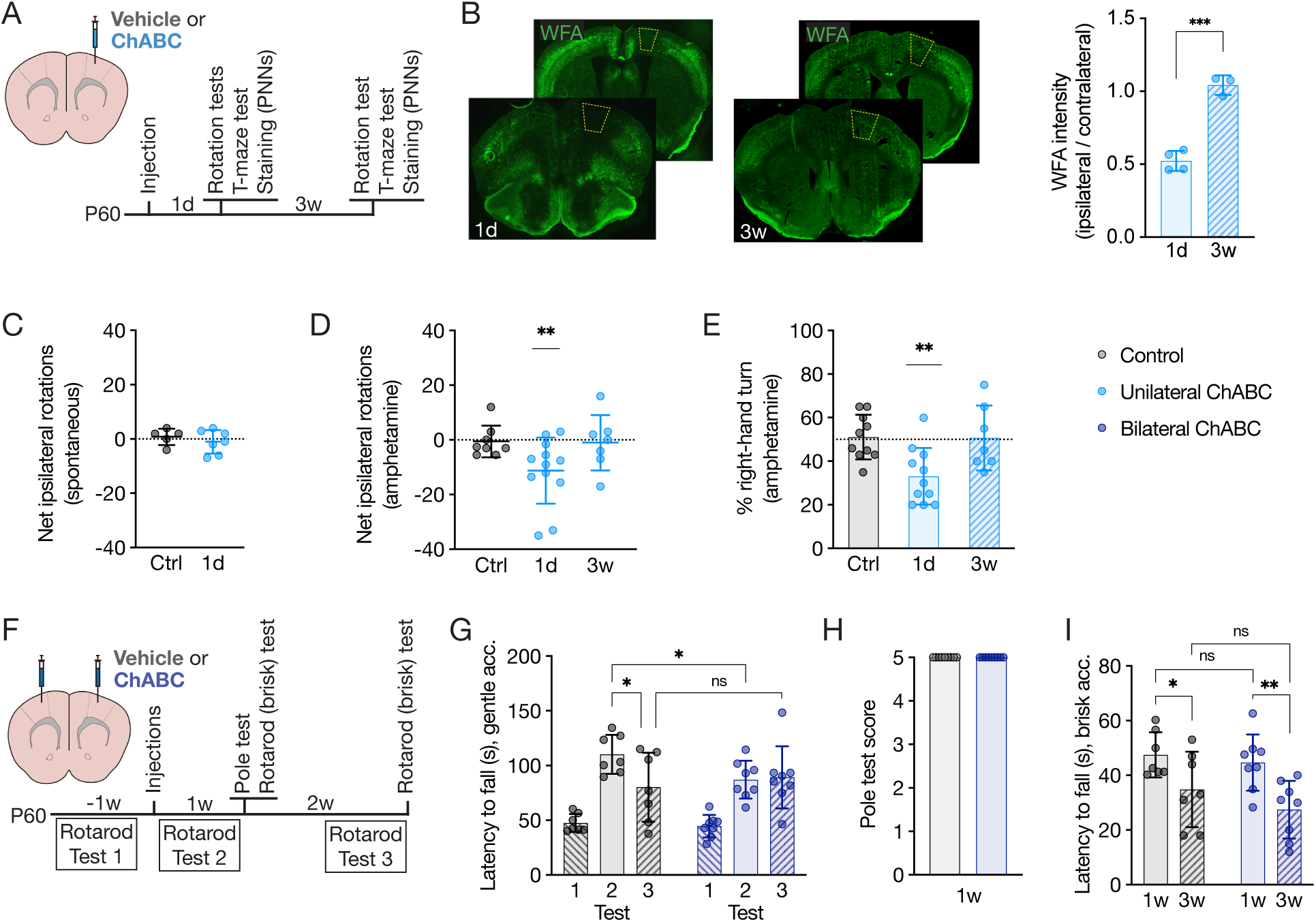
Chondroitinase ABC injections in M1 transiently alter motor function in healthy animals. **(A)** Timeline of experiments involving unilateral ChABC injection. **(B)** Representative images of WFA-stained sections of the anterior (front) and posterior (back) M1 cortex (yellow area) 1 day and 3 weeks after unilateral ChABC injection. Quantified WFA intensity of each section was normalized between ipsilateral and contralateral hemisphere. Unpaired *t*-test; n per group = 4 (unilateral ChABC, 1d) or 3 (unilateral ChABC, 3w). **(C)** Spontaneous rotations measured 1 day after injection. One-sample *t*-test compared to 0 net rotations; n per group = 5 (Control) or 7 (unilateral ChABC). **(D)** Amphetamine-induced rotations measured 1 day or 3 weeks after treatment. One-sample *t*-test compared 0 net rotations; n per group = 8 (Control), 12 (unilateral ChABC, 1d), or 7 (unilateral ChABC, 3w). **(E)** Amphetamine-induced T-maze alterations measured 1 day or 3 weeks after treatment. One-sample *t*-test compared to 50% right-hand turns; n per group = 10 (Control), 11 (unilateral ChABC, 1d), or 7 (unilateral ChABC, 3w). **(F)** Timeline of bilateral ChABC experiment. **(G)** Gentle accelerating rotarod performance reported as the latency to fall from the rod. 2-way ANOVA, uncorrected Fisher’s LSD; n per group = 7 (Control) or 8 (bilateral ChABC). Note that values are significantly higher (*P* < 0.0001) in Test 2 compared to Test 1 for both groups. **(H)** Pole test score 1 week after treatment. Unpaired *t*-test; n per group = 7 (Control) or 8 (bilateral ChABC). **(I)** Briskly accelerating rotarod performance reported as the latency to fall from the rod measured 1 week and 3 weeks after treatment. One-way ANOVA, Tukey’s Test; n per group = 7 (Control) or 8 (bilateral ChABC). All values ± SD; * *P* < 0.05, ** *P*<0.01, *** *P* < 0.001; ns, non-significant.

To determine the effect of ChABC treatment on symmetrical motor tasks, bilateral ChABC injections in M1 were performed, and the rotarod test, which necessitates symmetrical coordination and balance, was used to assess motor behavior (Fig. 1F). Following pre-screening sessions on a rotarod with gentle acceleration (Test 1) that also served as habituation, ChABC or vehicle solution was injected in both M1 hemispheres. One day later, animals were engaged in daily motor tests on the gently accelerating rotarod (Test 2) and, on the fifth day, motor performance was assessed with briskly accelerating rotarod and pole descent tests which engage both limbs. A significant yet small decrease in gentle acceleration rotarod performance for ChABC-injected mice was observed during Test 2 compared to control mice (Fig. 1G), which was most evident during the first and second days of testing after injection (Fig. S1C). However, at 1 wpi, all mice performed similarly on average in the pole test and the briskly accelerating rotarod (Figs. 1H and 1I). After a weeklong rest period, daily fixed-speed rotarod training (Test 3) showed no difference in motor performance between the two groups (Fig. 1G), culminating in no significant difference in either accelerating rotarod or pole test performance at 3 wpi (Figs. 1H and 1I). Both groups showed reduced accelerating rotarod performance at 3 wpi compared to 1 wpi (Fig. 1I). Together, these results suggest that bilateral ChABC injections impart very limited modifications on motor task learning that are inconsequential for motor performance.

### Midbrain lesions affect M1 PV cells

The amphetamine-dependent asymmetric motor behavior upon unilateral ChABC injection (Fig. 1C-E) suggests neuromodulation by dopamine (DA) given that amphetamine induces DA release, notably from midbrain TH neurons. Furthermore, the learning of motor tasks, such as the rotarod test, involves dopaminergic (DArgic) connections in M1 (33). This led us to characterize M1 PV cells in a PD model, where the physiopathology is characterized by a lack of DA. Unilateral lesions of the SNc were generated by acute 6-OHDA injections in the medial forebrain bundle (MFB) with an optimal dose of 2.5 µg that induced reliable motor deficits without severely compromising general health (Figs. S2A and S2B). At 2 wpi, lesions were confirmed anatomically by a loss of ∼80% TH^+^ cells in the SNc and ∼40% in the VTA (Fig. 2B), and behaviorally by spontaneous rotation behavior (Fig. 2C). After 3-4 wpi, mice have recovered from a post-operation lethargic state and are able to perform more complex motor tasks. At 5 wpi, lesioned mice showed strong behavioral deficits in the fixed-speed rotarod test, the pole test, and the beam test (Figs. 2D, 2E and 2F). Asymmetrical behavior was not observed with these tests, not even in the beam test for which paw slips were equal on either ipsilateral or contralateral sides (Fig. S2C).

**Figure 2.**
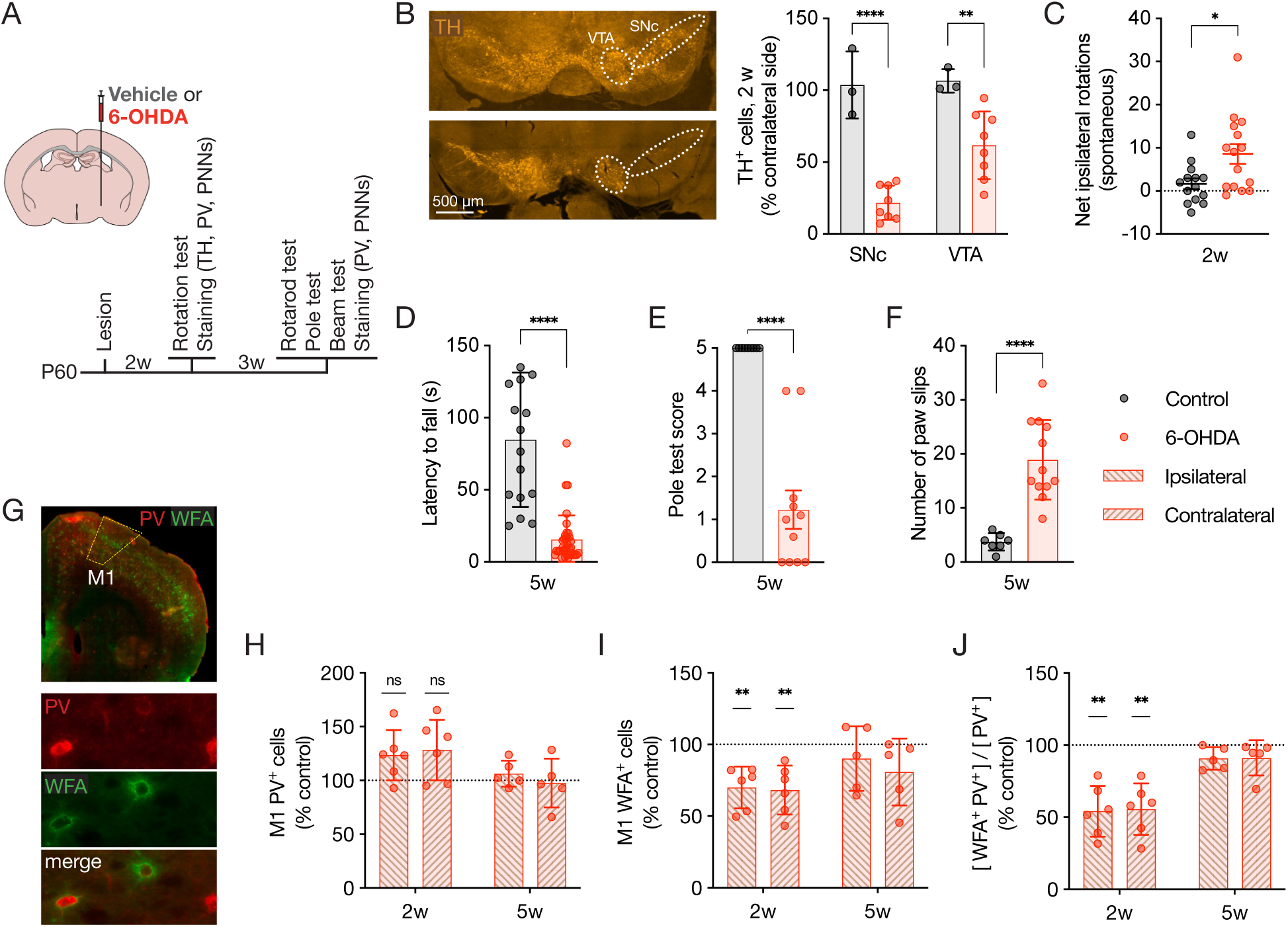
6-OHDA lesions generate reliable motor phenotypes and transiently alter M1 PV cells. **(A)** Timeline of the 6-OHDA lesion experiments. **(B)** Representative images and quantification of TH-stained sections of the Control (top) and 6-OHDA injected (bottom) SNc and VTA (white areas) 2 weeks after injection. Unpaired *t*-test; n per group = 6 (Control) or 8 (6-OHDA). **(C)** Spontaneous rotations measured 2 weeks after injection. One-sample *t*-test compared to 0 net rotations; n per group = 14 (Control) or 15 (6-OHDA). **(D)** Fixed-speed rotarod performance 5 weeks after treatment reported as the latency to fall from the rod. Unpaired *t*-test; n per group = 15 (Control) or 35 (6-OHDA). **(E)** Pole test score 5 weeks after treatment. Unpaired *t*-test, n per group = 9 (Control) or 11 (6-OHDA). **(F)** Irregular beam performance 5 weeks after treatment reported as the number of paw slips. Unpaired *t*-test; n per group = 7 (Control) or 12 (6-OHDA). **(G)** Representative images of PV and WFA staining in M1 cortex (white area). **(H-J)** Quantification of PV^+^ cells (H), WFA^+^ cells (I) and double-stained PV cells (J) in either M1 hemisphere of lesioned mice at 2 and 5 weeks, normalized to control levels. One-sample *t*-test compared to 100%; n per group = 6 (2w) or 5 (5w). All values ± SD; * *P* < 0.05, ** *P* < 0.01, **** *P* < 0.0001; ns, non-significant.

To investigate the effect of 6-OHDA lesions on PV cells in M1, double staining for PV and PNNs (PV^+^ and WFA^+^ cells) on sections encompassing both anteroposterior and mediolateral M1 were analyzed at 2 and 5 wpi (Fig. 2G). During the lethargic state at 2 wpi, no significant impact on the number of PV cells was observed in either hemisphere (Fig. 2H). In contrast, the number of WFA^+^ cells decreased across all M1 layers in both ipsilateral and contralateral hemispheres, despite the lesion being unilateral (Fig. 2I). This effect was particularly robust on PV^+^ cells (Fig. 2J), indicating a more pronounced disruption of PNNs of PV cells. Because PNNs can be sensitive to oxidative stress, we controlled for a potential global oxidizing effect of 6-OHDA. At 2 wpi, the numbers of PV^+^ cells, WFA^+^ cells, and double-stained cells were found to be unchanged in V1, a cortical area not directly involved in motor pathways (Figs. S2D-G). Finally, although motor behavior is greatly impacted at 5 wpi, PNN levels are recovered in both hemispheres (Figs. 2I and 2J), indicating that this morphological effect on M1 PV cells is transient.

### Motor performance of lesioned animals is improved by the disruption of M1 PNNs

Given the potential for PNN removal to induce adult plasticity and ameliorate behavioral outcomes in stroke animal models (26, 27), we evaluated the effects of ChABC injections in M1 on the motor behavior of 6-OHDA-lesioned animals. Since unilateral ChABC injections induce lateralized motor behavior in healthy animals, we initially focused on bilateral ChABC injections. A novel cohort of animals was injected with 6-OHDA and, after 4 wpi, a three-day “pre-screen” fixed-speed rotarod assessment (“0w”) was conducted for habituation and to obtain a baseline performance (Fig. 3A). The lesioned animals were split homogeneously into two groups based on pre-screen performance, and at 5 wpi, one group received bilateral injections of ChABC in M1 while the other received a vehicle solution. Previous research has underscored the pivotal role of motor training in facilitating successful recovery of motor disorders (34, 35). To provide motor stimulation, animals were then housed in cages enriched with a running wheel, shelter, and a tube for the rest of experiment. After a 24-hour recovery period, animals underwent daily motor training sessions consisting of three trials on the rotarod and one trial on the pole test, for five consecutive days (“1w”) to provide additional motor stimulation. After a one-week rest period, subsequent testing sessions were conducted (“3w”). Despite PNN recovery, lesioned animals without ChABC injection displayed consistent poorly performing behavior over the three sessions, with no improvement in rotarod nor pole performance after 1w and 3w training (Figs. 3B and 3C). The non-lesioned control group performed significantly better after the 0w pre-screen session, suggesting a motor learning effect in the rotarod task (Fig. 3B). Interestingly, the lesioned group injected with ChABC also showed improved performance during the 1w and 3w testing sessions compared to the 0w pre-screen (Fig. 3B). Furthermore, this group showed improved performance on the pole test between 0w, 1w and 3w, indicating persistent improvement in motor performance (Fig. 3C).

**Figure 3.**
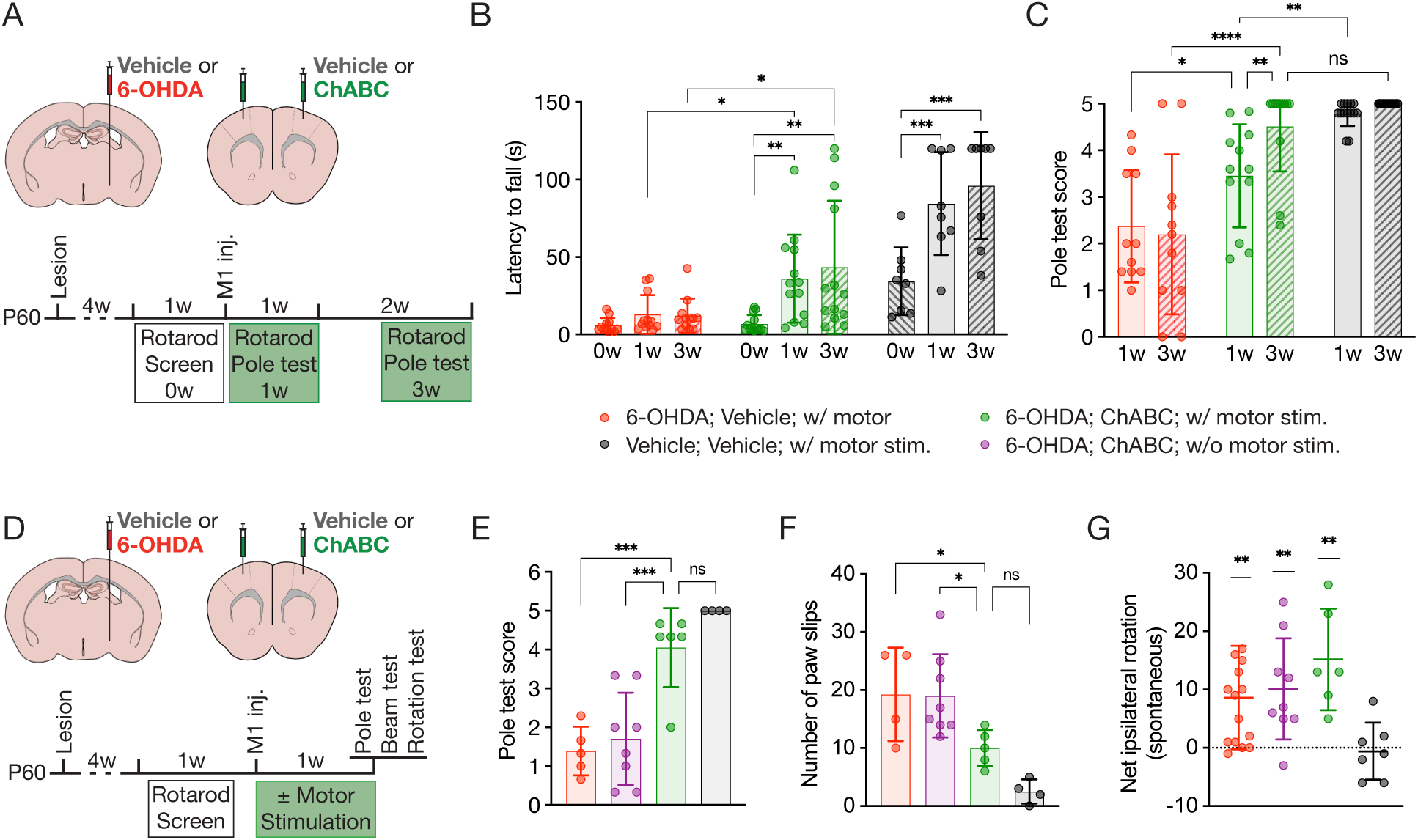
Bilateral ChABC injections coupled with motor stimulation improve motor control in lesioned mice. **(A)** Timeline of the experiments of ChABC treatment and motor stimulation for behavior rescue in lesioned mice. **(B)** Fixed-speed rotarod performance reported as time to fall from the rod in animals with motor stimulation (i.e., motor training and housing enrichment). 2-way ANOVA, uncorrected Fisher’s LSD; n per group = 8 (Control), 12 (6-OHDA; Vehicle), or 13 (6-OHDA; ChABC). Note that Control values are significantly higher (*P* < 0.01 or 0.001) than 6-OHDA lesioned mice (w/o or w/ ChABC) for each session. **(C)** Pole test score 1 and 3 weeks after ChABC surgery. 2-way ANOVA, uncorrected Fisher’s LSD; n per group = 14 (Control), 11 (6-OHDA; Vehicle), or 12 (6-OHDA; ChABC). Note that Control values are significantly higher (*P* < 0.0001) than 6-OHDA lesioned mice (w/o ChABC) for each session. **(D)** Timeline for assessing the effect of motor stimulation (i.e., motor training and housing enrichment) in ChABC-dependent behavioral rescue of lesioned mice. **(E)** Pole test score 1 week after ChABC treatment. One-way ANOVA, Holm-Šídák’s test; n per group = 4 (Control w/ stimulation), 4 (6-OHDA; Vehicle; w/ stimulation), 8 (6-OHDA; ChABC; w/o stimulation) or 6 (6-OHDA; ChABC; w/ stimulation). Note that Control values are significantly higher (*P* < 0.0001) than for 6-OHDA lesioned mice treated w/ Vehicle; w/ stimulation or w/ ChABC; w/o stimulation. **(F)** Irregular beam balance performance reported as number of paw slips, 1 week after ChABC surgery with or without motor training. One-way ANOVA, Tukey’s Test; n per group = 4 (Control; w/ stimulation), 4 (6-OHDA; Vehicle; w/ stimulation), 8 (6-OHDA; ChABC; w/o stimulation) or 6 (6-OHDA; ChABC; w/ stimulation). **(G)** Spontaneous rotation 1 week after ChABC surgery with or without motor training. One-sample *t*-test compared to 0 net rotations; n per group = 7 (Control w/ stimulation), 14 (6-OHDA; Vehicle w/ stimulation), 9 (6-OHDA; ChABC w/o stimulation) or 6 (6-OHDA; ChABC; w/ stimulation). All values ± SD; * *P* < 0.05, ** *P* < 0.01, *** *P* < 0.001, **** *P* < 0.0001; ns, non-significant.

The motor improvements associated with bilateral ChABC injection could be either solely dependent on the molecular effect of ChABC or contingent on the motor stimulation provided by enriched housing and daily testing sessions. To distinguish between these possibilities, a novel cohort of lesioned animals was further divided into groups receiving bilateral ChABC injections coupled either with motor stimulation via enriched cages and daily rotarod test sessions or without such stimulation (Fig. 3D). To avoid confounding effects from motor learning, all animals were subjected to novel tests in order to evaluate motor performance. The ChABC treated animals receiving motor stimulation performed significantly better than non-stimulated animals in both the pole test (Fig. 3E) and the irregular beam test (Fig. 3F). In the pole test, lesioned mice also displayed akinetic behavior that was only reversed when ChABC was combined with motor stimulation (Fig. S3A). Thus, the rescue effect of bilateral PNN removal in M1 clearly depends on motor stimulation. Daily motor stimulation was not found to affect the motor performance of control animals nor of lesioned animals untreated with ChABC (Fig. S3B and S3C).

To determine whether treatment of lesioned animals with motor stimulation and M1 PNN removal can overcome asymmetrical motor behavior, animals were also assessed for spontaneous rotation behavior (Fig. 3G). Significant ipsilateral rotations were still observed in the ChABC rescue groups regardless of motor stimulation, suggesting that the asymmetrical phenotype is not affected by ChABC treatment. Given that unilateral ChABC injections in healthy mice result in amphetamine-induced contralateral rotational behavior (Fig. 1D), we measured the effect of unilateral ChABC injections on the rotational behavior of lesioned animals to determine the possibility of additive or subtractive mechanisms. Lesioned animals were injected unilaterally in M1 with ChABC, either ipsilateral or contralateral to the lesioned hemisphere, and spontaneous rotational behavior was assessed 1d later (Fig. 4A). Rotation behavior was annulled by the ipsilateral ChABC injections yet maintained by the contralateral injections (Fig. 4B). Given that ipsilateral-injected animals displayed no rotational behavior, we tested whether unilateral PNN removal in ipsilateral M1 also provides motor performance rescue similar to bilateral injections. A unilateral ChABC injected cohort was also submitted to the motor stimulation paradigm with enriched housing and rotarod tests (Fig. 4C), similar to the bilaterally injected cohorts (Figs. 3A and 3B), yet no motor performance improvement was observed (Fig. 4D). Thus, PNN removal is required in both M1 hemispheres for the rescue of symmetrical motor phenotypes in lesioned animals.

**Figure 4.**
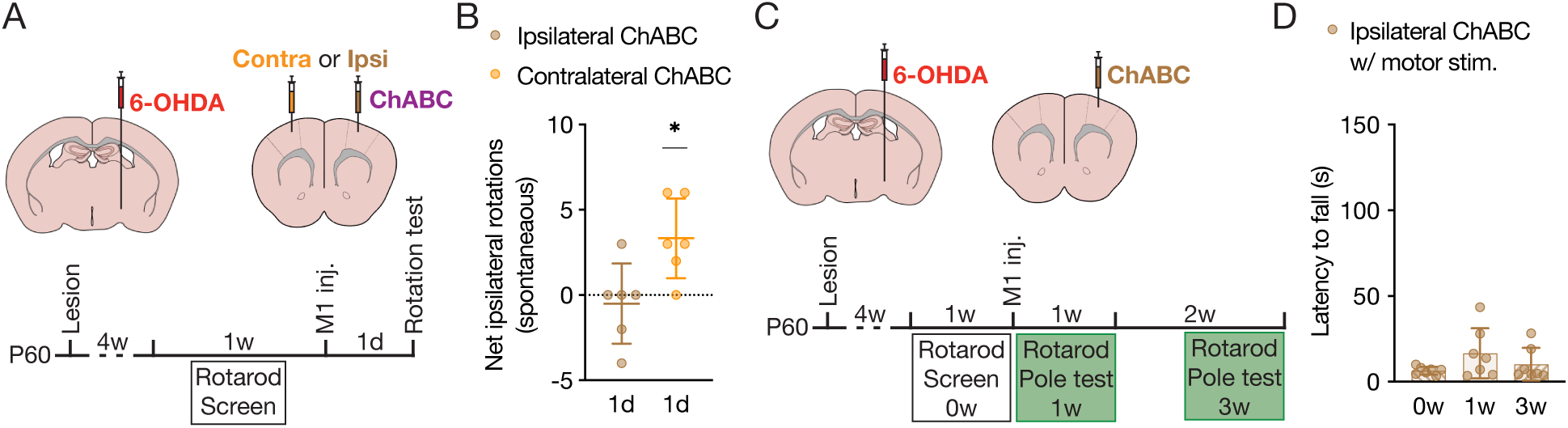
Unilateral ChABC injections coupled with motor stimulation do not improve motor control in lesioned mice. **(A)** Timeline of experiments on lesioned mice with unilateral ChABC experiment. **(B)** Spontaneous rotations measured 1 day after ChABC treatment. One-sample *t*-test compared to 0 net rotations; n per group = 6. **(C)** Timeline of unilateral ChABC treatment and motor tests for behavior rescue. **(D)** Fixed-speed rotarod performance reported as time to fall from the rod for lesioned mice treated with an ipsilateral M1 ChABC injection. One-way ANOVA, Tukey’s Test; n = 7. All values ± SD; * *P* < 0.05.

## Discussion

In the mammalian cerebral cortex, mature PV cells are enwrapped by PNNs maintaining a non-plastic state by altering the extracellular microenvironment to provide molecular brakes and synapse stabilization. In V1, PNN digestion by ChABC impacts PV cell activity by increasing the recruitment of thalamocortical afferents, which affects state-dependent feed-forward inhibition (21), and by reducing PV cell intrinsic activity, which results in decreased stability of the excitatory-inhibitory (E/I) balance (36). A shift in E/I balance towards excitation would result in increased principal neuron activity. Based on these premises, a first hypothesis explored in this work was that the unilateral degradation of PNNs in M1 would enhance upper motoneuron (UMN) activity resulting in spontaneous rotations contralateral to the ChABC-injected side. While this was not the case, an increase in contralateral rotations was nevertheless obtained through amphetamine-induced DA secretion. The discrepancy between spontaneous and amphetamine-induced behavior suggests a subtle effect of ChABC that is compensated in normal conditions. As amphetamine treatment leads to increased DA release, the loss of PNNs might perturb M1 PV cell-mediated DA response affecting UMN activity. For example, the loss of PNNs in the anterior cingulate cortex enhanced DA-mediated local high-frequency network activity (37). A possible consequence of the loss of PNNs is to limit the ability of PV cells to maintain high firing rates (38), which may alter cortical E/I balance only in the context of increased motor activity. However, we cannot distinguish whether the effects are due to DA acting directly in M1 following its liberation from VTA terminals or indirectly, and more classically, through the SNc-striatum-thalamus-M1 pathway (39–41). Regardless, this result suggests an interplay between PNN assembly in M1 and the activity of DArgic pathways, with motor outcomes. Indeed, asymmetry in motor behavior was lost 3 weeks later after PNN reassembly.

To investigate the possible interplay with DArgic pathways, PNN assembly in M1 was evaluated following a unilateral 6-OHDA injection in the MFB, leading to ∼40% and ∼80% loss of TH-expressing cells in the VTA and SNc, respectively. Two weeks after lesioning, mice showed spontaneous ipsilateral rotations, in keeping with the expected loss of mDA afferent activity in the striatum and, directly or indirectly, in M1. Yet, surprisingly, 50% of M1 PV cells had lost their PNNs not only on ipsilateral side, but also on the contralateral side. Five weeks after lesioning, the number of PNN-enwrapped PV cells was restored in both hemispheres. This unexpected bilateral PNN loss cannot be explained by a general oxidative stress induced by 6-OHDA as PNNs were found to be intact in V1. Bilateral downregulation of PNNs have been observed in the lateral vestibular nucleus following unilateral peripheral vestibular lesion (42), and in the motor cortex following a unilateral stroke (26), suggesting that bilateral M1 regulation may be important for compensation. This bilateral response to the loss of unilateral mDA cells and projections may result from cortico-cortical interactions which can be direct through the corpus callosum involving principal neurons or through a subpopulation of PV cells with transcortical projection (CC-PV cells) accounting for about 40% of the total PV cell population (43–45). Inter-hemispheric connections in motor function were implicated in a traumatic brain injury model in which M1 unilateral ChABC treatment upregulated neuronal activity not only in the ipsilateral cortex but also in the contralateral cortex (46). Another possibility is the intra-telencephalic (IT) pathway projecting to contralateral cortex or striatum (47, 48). Finally, reduced motor activity might also be contributing to this bilateral loss of M1 PNNs. As reductions in network activity are sufficient to induce PNN regression in V1 (49), the decreased locomotion may have a similar effect in M1 when animals are very lethargic during the ∼3-week recovery phase following 6-OHDA injection.

As anticipated, mice continue to exhibit net ipsilateral rotations 5 weeks after lesioning, as the unilateral loss of mDA neurons is permanent. Behaviors requiring bilateral coordination, such as the rotarod, pole, and beam tests, are also greatly impacted even though animals have recovered from their state of lethargy. The transient reduction in PNNs at 2 weeks during this initial state, suggests transient M1 circuit plasticity given that PNN-negative PV cells are prone to enhanced plasticity (38). It has been shown that 2 weeks after 6-OHDA SNc lesions, M1 circuitry preserves E/I balance yet manifests altered intracortical inhibitory synaptic drive, with PV cells showing impaired excitability in response to high-frequency activity (50, 51). At 5 weeks post-lesion with full recovery of M1 PNNs, we hypothesized that motor performance remains limited by maladapted M1 circuitry. PNN removal by ChABC treatment in V1 is able to cure experimental amblyopia in a binocular environment (binocular training). Based on this analogy, we tested whether bilateral M1 PNN degradation 5 weeks post-lesion in association with motor stimulation (enriched housing coupled with motor training) could improve bilateral motor behaviors. This was the case, with the additional finding that motor stimulation was mandatory but did not have to be specific given that rotarod training enhances performance in both pole and beam tests. Nevertheless, spontaneous ipsilateral rotations were maintained in all conditions, confirming that turning behavior can be separated from behaviors requiring bilateral coordination, even if they were impacted by the unilateral lesion.

The distinction between turning behavior and bilateral motor coordination is evident when ChABC is injected either ipsilateral or contralateral to the lesioned hemisphere at 5 weeks post-lesion when M1 PNN levels are restored. One day after treatment, rotations were abolished only by an ipsilateral ChABC injection in M1. Unilateral PNN removal may increase UMN activity to induce contralateral turning and counterbalance the effects of the 6-OHDA lesion. However, such an explanation runs counter to unilateral ChABC injections not inducing spontaneous contralateral rotations in non-lesioned mice. In lesioned mice, perhaps, the decrease in PV cell inhibitory activity resulting from PNN removal is additive with that due to the loss of GABA release resulting from the decrease in local DA release for UMN activation (33, 40, 41). Added together these two events would decrease spontaneous contralateral turning and compensate for the loss of mDA neuron activity. However, this unilateral rescue effect is transient and ipsilateral ChABC treatment does not lead to improved motor performance even when associated with motor stimulation. An improvement in the bilateral motor behavior of lesioned mice requires bilateral PNN removal in M1.

Drawing parallels between motor and visual cortex plasticity, it is evident that similar principles govern neural remodeling and functional recovery across different brain regions and sensory modalities. Enhanced plasticity coupled with appropriate cortical stimulation can help restore partial function after an insult. The transient nature of PNN disruption observed after mDA lesions echoes the dynamic nature of cortical plasticity in response to healthy or pathological stimuli. Leveraging our understanding of neural plasticity mechanisms across various neurological conditions, including PD, stroke, and visual impairment, holds promise for the development of targeted therapeutic interventions aimed at promoting functional recovery and improving quality of life.

Although electrophysiology studies suggest 6-OHDA lesions do not greatly impact M1 PV cell activity at either 2 weeks (51) or 3 weeks (31, 32) after lesioning, we find bilateral PNN disruption of M1 PV cells at 2 weeks. Circuit function is clearly being altered in response to reduced mDA activity and/or lethargic behavior, leading to long-lasting motor consequences and permanent circuit dysfunction as M1 PNNs recover. Remarkable functional improvement can be unlocked by PNN removal (Fig. 5). This finding underscores the prolonged repercussions of acute lesions and the importance of choosing pertinent timepoints for molecular, physiological and behavioral analyses. It will be of great interest to examine the impact and therapeutic potential of M1 PNNs in chronic PD animal models involving synucleinopathies with progressive loss of mDA activity.

**Figure 5.**
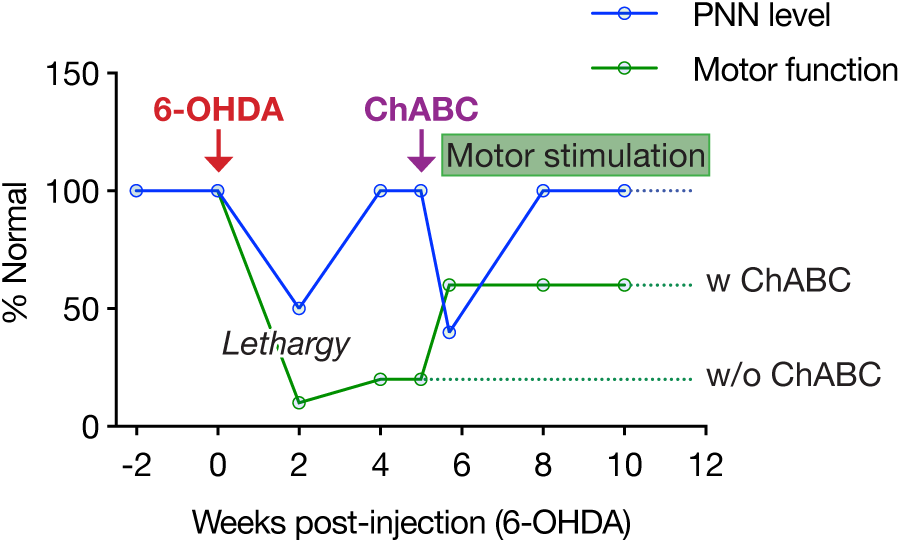
Schematic graph of the kinetics of the lesion-dependent and ChABC-dependent changes in PNN levels and motor behavior.

## Acknowledgements

We would like to thank Pierre-Alexandre Curty for technical assistance. Funding was provided by the Neuroglia Fund for supporting research costs and salaries.

---

**Figure S1 (related to Figure 1).**
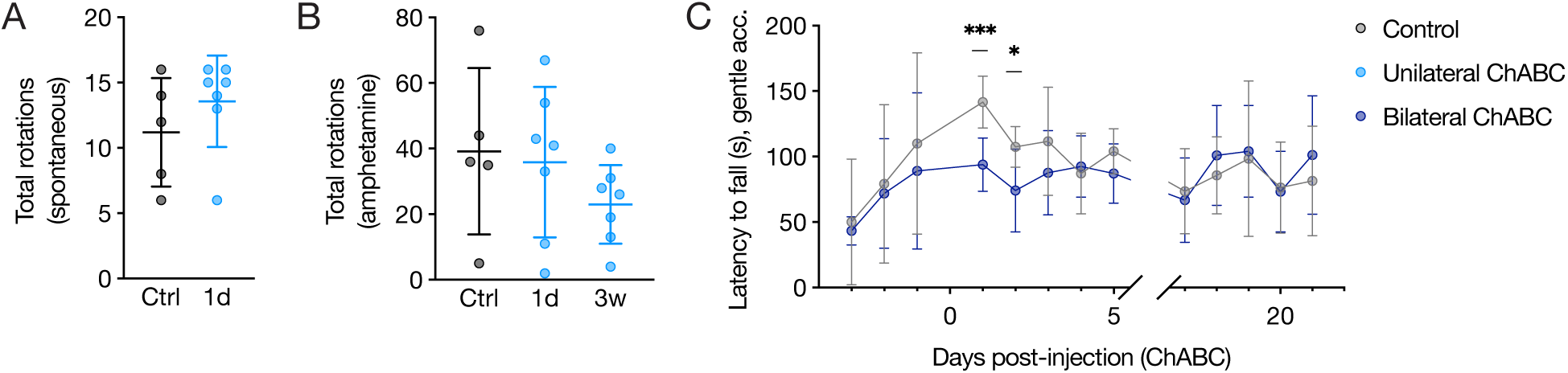
Rotation behavior and daily rotarod performance of mice with M1 ChABC injections. **(A)** No significant changes in total spontaneous rotations measured 1 day after M1 ChABC injection compared to control injection. Unpaired *t*-test; n per group = 5 (Control) or 7 (unilateral ChABC, 1d). **(B)** No significant changes in total amphetamine-induced rotations measured 1 day or 3 weeks after M1 ChABC injection compared to control injection. One-way ANOVA; n per group = 5 (Control) or 7 (unilateral ChABC, 1d and 3w). **(C)** Daily recordings of rotarod performance of animals with or without M1 ChABC injections. Gentle accelerating rotarod performance reported as the latency to fall from the rod. 2-way ANOVA, uncorrected Fisher’s LSD; n per group = 7 (Control) or 8 (bilateral ChABC). All values ± SD; * *P* < 0.05, *** *P* < 0.001.

**Figure S2 (Related to Figure 2).**
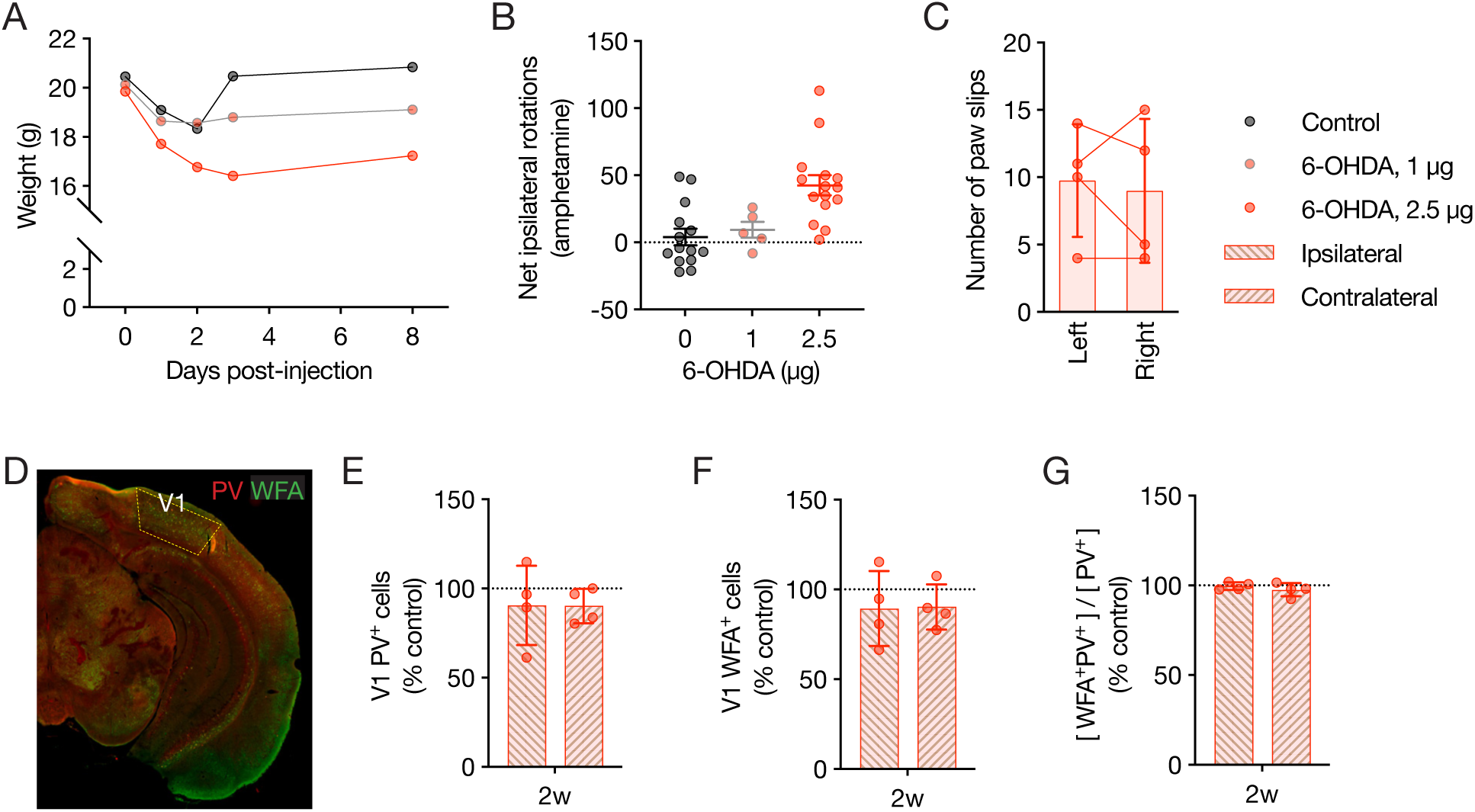
Characterization of 6-OHDA dose on animal physiology and V1 PV cells. **(A)** Weight loss progression following injection of vehicle or 6-OHDA (1 or 2.5 µg). n per group = 8 (Control), 6 (1 µg 6-OHDA), or 7 (2.5 µg 6-OHDA). **(B)** Amphetamine-induced rotations in Control and lesioned animals. One-sample *t*-test compared to 0 net rotations; n per group = 15 (Control), 5 (1 µg 6-OHDA), or 15 (2.5 µg 6-OHDA). **(C)** Irregular beam test performance 5 weeks after lesioning reported as number of paw slips comparing slips on either left or right side for each animal. Paired *t*-test; n per group = 4. **(D)** Representative image of PV and WFA staining in V1 (yellow area). **(E-G)** Quantification of PV^+^ cells (E), WFA^+^ cells (F) and doubled stained PV cells (G) in either V1 hemisphere of lesioned mice 2 weeks after treatment, normalized to control levels. One-sample *t*-test compared to 100%; n = 4. All values ± SD; * *P* < 0.05, ** *P* < 0.01, *** *P* < 0.001; ns, non-significant.

**Figure S3 (Related to Figure 3).**
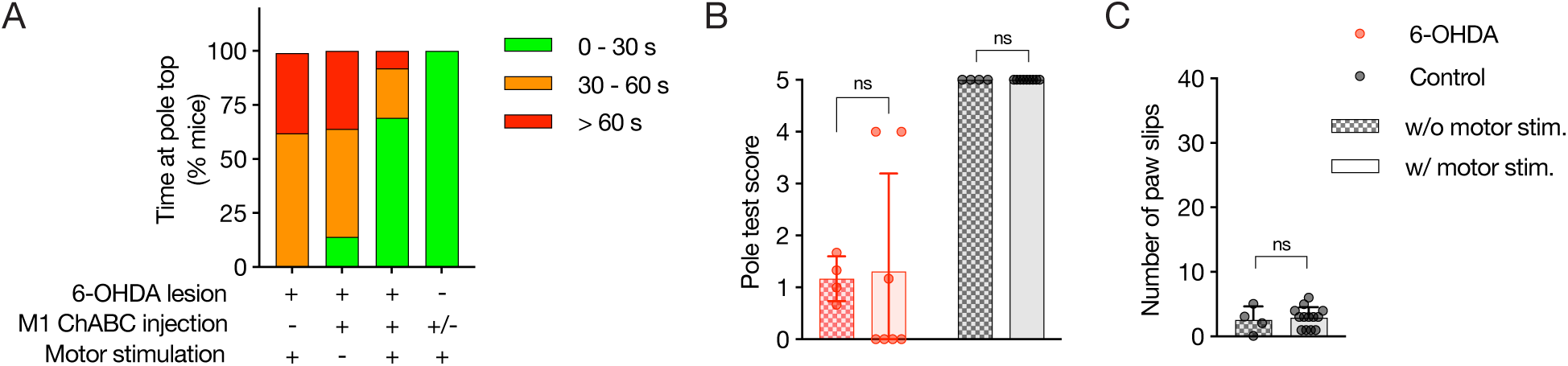
Effect of motor stimulation on lesioned and non-lesioned animals. **(A)** Akinesia in the pole test evaluated by the time spent at the top of the pole, with mice grouped by time brackets. **(B)** Pole test score 1 week after ChABC treatment with or without motor stimulation. Unpaired *t*-test; n per group = 4 (6-OHDA w/o stimulation), 7 (6-OHDA w/ stimulation), 4 (Control w/o stimulation), or 9 (Control w/ stimulation). **(C)** Irregular beam test performance reported as number of paw slips, 1 week after ChABC treatment with or without motor training. Unpaired *t*-test; n per group = 4 (Control w/o stimulation), or 13 (Control w/ stimulation). All values ± SD; ns, non-significant.

